# Sensing Plant Physiology and Environmental Stress by Automatically Tracking F_j_ and F_i_ Features in PSII Chlorophyll Fluorescence Induction

**DOI:** 10.1101/362939

**Authors:** Qian Xia, Jinglu Tan, Shengyang Cheng, Yongnian Jiang, Ya Guo

## Abstract

Following a step excitation, chlorophyll fluorescence (ChlF) from photosystem II of a dark-adapted plant leaf exhibits the well-known OJIP pattern. The OJIP induction has been widely applied in plant science, agriculture engineering, and environmental engineering. While the J and I phases are related to transitions of photochemical reaction redox states, characteristic fluorescence intensities for the two phases (F_j_ and F_i_) are often treated as fixed time points in routine measurement and thus do not account for variations in plant and experimental conditions, which (1) neglects the time differences, potentially useful information for characterizing plant status and environmental factors, and (2) leads to errors in measured F_j_ and F_i_ values. In this work, a method for consistent measurement of F_j_ and F_i_ was developed through polynomial fitting and curvature analysis. The method measures the curvatures in the OJIP curve and automatically tracks the characteristic transition points under variable sample and experimental conditions. Experiments were carried out to demonstrate the concept and classification capabilities of the developed method. This research established a new framework to analyze ChlF and enhanced the applications of ChlF.

## 1. Introduction

Light absorbed by plant photosystem II (PSII) in the photosynthetic process has three subsequent pathways: photochemical reactions, heat and chlorophyll fluorescence (ChlF) (Goltsev et al., 2003; Krause and Weis, 1991; Lavergene and Trissl, 1995; Stirbet et al., 1998; Vredenberg, 2004; Taiz and Zeiger, 2006). Since the emission of ChlF competes for light energy with the other two pathways (Lubitz et al., 2008), almost all the changes in photosynthesis can be reflected by PSII ChlF (Zhu et al., 2005; Rodriguez & Greenbaum, 2009). ChlF measurement is thus a reliable method to study plant physiology and environment factors that influence photosynthesis (Mohammed et al., 1995; Maxwell & Johnson, 2000; Coombs & Long, 2014). In addition, ChlF is a fast, noninvasive, simple, and intuitive way to represent the changes in photosynthetic activities (Kootenet et al., 1990; Mathur et al., 2011).

When a step illumination is applied, the intensity of PSII ChlF from a dark-adapted plant leaf changes with time in a unique pattern of the commonly-labeled O-J-I-P phases, which is referred to as the ChlF induction curve or OJIP curve. The OJIP curve contains abundant information about the process of photosynthesis: the absorption and conversion of light energy, the transfer and distribution of energy, the state of the reaction centers, the activities of the PSII donors and the receptors, plastoquinone (PQ) pool size and the activities, excess light energy and its dissipation, photosynthetic light suppression, and light damage. ChlF induction has been extensively used in the literature to analyze plant photosynthesis and physiological conditions (Ogaya et al., 2011; Schansker et al., 2014; Guo & Tan, 2015; Guo et al., 2015).

There are four important points on an OJIP curve. O reflects the initial fluorescence when a leaf is exposed to light after dark adaptation. J indicates accumulation of plastoquinone A (Q_A_^−^) (Strasser et al., 1995). I is related to the heterogeneity of the PQ pool (Strasser et al., 1995; Jee, 1995). P shows the maximum value of fluorescence. F_j_ and F_i_ represent the ChlF intensity during the J and I phases of the ChlF kinetics, respectively. Obviously, these time points depend on the photochemical reaction kinetics, which implies that differences in plant physiological and experimental conditions may lead to different times of occurrence for these phases. J and I are generally defined as the first and the second inflection points or intermediary peaks on the ChlF induction curve, respectively (see Figure 1). Different plant species, light intensity, temperature, salinity, and drought may affect the plant physiological status (D’Ambrosio et al., 2006; Koyro, 2006; Ruban & Belgio, 2014; Guo & Tan, 2015) and thus the times of occurrence of these transitions.

**Fig. 1.**
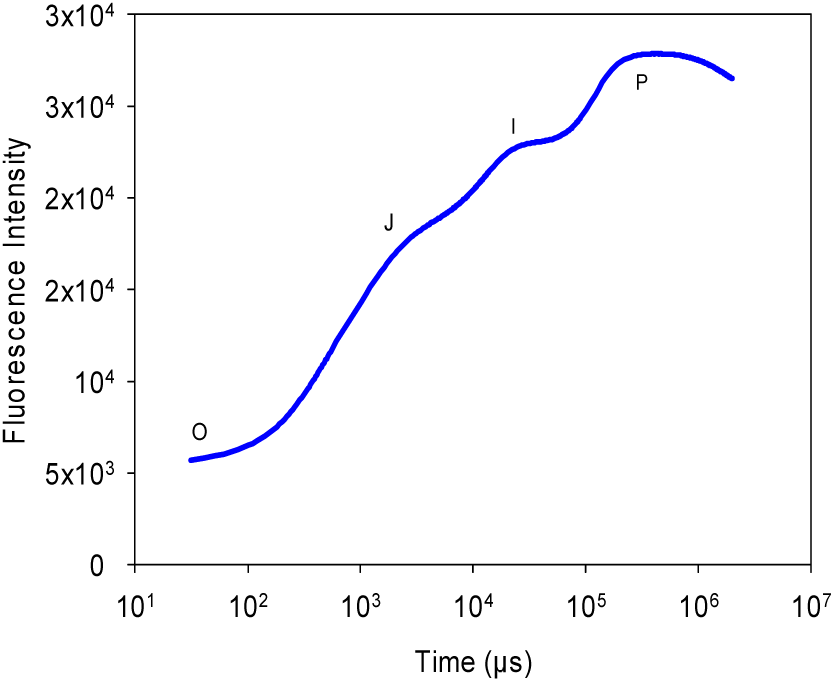
A typical ChlF induction or OJIP curve

The direct ChlF induction parameters include F_o_, F_j_, F_i_, and F_m_. Many other ChlF parameters can be computed from these four. F_o_ is defined as the initial fluorescence and thus it can be measured at a fixed time after the initiation of excitation. F_m_ represents the maximum fluorescence in the entire ChlF induction curve and thus the exact time of F_m_ measurement is not critical. J and I, however, are transient phases between O and P. They may occur at different times depending on the plant species, illumination, and growth environments. Currently, all commercial ChlF instruments measured F_j_ and F_i_ at a fixed predefined time, although it is manually adjustable. The instruments cannot automatically adjust the characteristic time and track the F_j_ and F_i_ points. Although users may configure the predefined time through operating the instrument menu, but it is boring and time consuming to set it for each measurement and ChlF features were usually read according to a fixed time. This has several obvious drawbacks: (1) It neglects the time differences, which may be useful information for characterizing plant status and environmental factors because these times of occurrence reflect the reaction rates, and (2) It leads to errors in measured F_j_ and F_i_ values because a fixed time is inappropriate for all plant and experimental conditions. This may limit the ChlF usefulness and results in discrepancies in interpretation of ChlF kinetics.

This work was aimed at developing a method to determine the J and I characteristic times adaptively and consistently when plant and experimental conditions vary. Based on the common interpretations of the ChlF induction kinetics, these times were determined according to the curvature changes on the OJIP curve. Least-squares polynomial fitting was used to filter experimental data. Comparisons were made between the proposed method and the traditional method. Applications were used to validate the usefulness of the proposed method.

## 2. Method Development

A typical OJIP induction curve is shown in Figure 1. Since J and I are inflection points in ChlF intensity resulting from changes in forward (downstream) reactions (Strasser et al., 1995; Stirbet and Govindjee, 1992), they are simply points of curvature changes in the induction curve. Consequently, they can be conveniently and consistently located by finding the local curvature maxima on the induction curve. Because the induction curve is commonly observed in semi-log scale, curvature is computed based on logarithm of time.

The curvature of a curve can be easily found by computing numerical differences, but numerical differences of a measured curve easily suffer from noise, resulting in difficulty in finding the true local maxima of curvature. To eliminate the influence of noise, the induction curve may be first fitted with a spline or polynomial of the form.

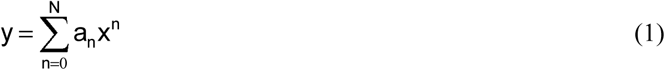

where *y* is ChlF, *x* is the logarithm of time (as the induction curve is usually presented in semi-log scale to reveal J and I), N is the order of the fitted polynomial, *a*_*n*_ (*n* = 0 … *N*) are coefficients, and *n* is an integer.

From Eq. (1), the first derivative of *y* with respect to *x* can be computed as:

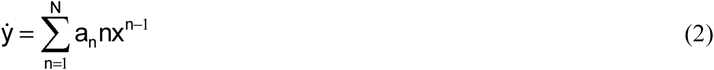

and the second order derivative *ÿ* is:

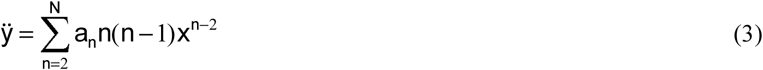

Curvature *k* is computed as:

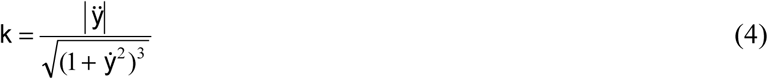

In this work, *N* was experimentally determined. When *N* increases, the fitting error for Eq. (1) will decrease. The N-value where the fitting error starts to level off was selected as the desired value and it was 15 in this work. The polynomial fitting is used to capture the main variations in ChlF and thus to eliminate noise. As a result, the exact order of the polynomial will not affect the analysis and the computed derivatives significantly. When the curvature *k* is obtained, it is easy to determine the times of J and I based on the *y* (OJIP) pattern and the *k* pattern. Obviously, either *y* or *k* may be rescaled or shifted for easy identification of the maxima without affecting the final outcomes.

## 3. Experiments

### 3.1 Plant Samples

Three types of tree leaves (*Cercis chinensis, Pittosporum tobira* and *Elaeocarpus glabripetalus merr*) and two types of vegetable leaves (spinach and lettuce) were used in the experiments.

The tree leaves were collected from the campus of Jiangnan University (Wuxi, China). The leaves were picked in the morning between 6 am and 7 am in July, when the environment temperature was around 28°C. The picked tree leaves were from the middle of the tree canopy on the south side. The two types of vegetables were acquired from a local farmers’ market in an early morning in July with an environmental temperature at approximately 28°C. The spinach and lettuce leaves are intact and fresh. The spinach had roots but the lettuce roots had been cut. The samples were transported to the laboratory for experiments. The laboratory temperature was at 25°C.

In order to reduce evaporation and influence of water status on ChlF emission, all the sample leaves were floated in water for at least half a day. Before ChlF measurement, the leaves were dark-adapted for at least 30 minutes in dark-adaption clips. ChlF was measured with a FluorPen PSI (Photon Systems Instruments, Czech Republic) by selecting its OJIP protocol.

### 3.2 Experimental Design and Data Analysis

In this work, four sets of experiments were designed to demonstrate the concept and usefulness of the proposed method. Twenty-nine leaves were measured for each variety of plant in the first three sets of experiments. For the fourth set of experiments, fourteen leaves of each plant type were measured.

The first set of experiments involved all five different types of leaves and was used to show the differences in the times of occurrence and F_j_ and F_i_ values from different plants obtained by the developed method or the default ChlF meter protocol. The illumination light intensity was set as 3,000 μmol photons m^−2^s^−1^.

The second set of experiments included four different leaves (spinach, *Cercis chinensis, Pittosporum tobira* and *Elaeocarpus glabripetalus merr*) and was used to compare the differences in the times of occurrence and F_j_ and F_i_ values under varied light intensities obtained by the developed method or by the default ChlF meter protocol. Three levels of light intensity were used: 750 μmol photons m^−2^s^−1^, 1,500 μmol photons m^−2^s^−1^, and 3,000 μmol photons m^−2^s^−1^. Half an hour dark-adaptation was applied in between measurements.

The third set of experiments involved the three types of tree leaves and was used to compare the differences in the times of occurrence and F_j_ and F_i_ values in various temperatures obtained by the developed method or by the default ChlF meter protocol. Each leaf was cut into two halves along the central vein. The leaf halves were considered to be in similar physiological status. One half was placed in water at 23°C and the other in water at 33°C. They were all processed at the same time. The illumination light intensity was set at 3,000 μmol photons m^−2^s^−1^.

The fourth set of experiments involved all five plant types and was used to compare the differences in the time of occurrence and F_j_ and F_i_ value under different levels of detachment stress obtained by the developed method or by the default ChlF meter protocol. All the leaves in this set of experiments were kept in 2°C. Wet paper towels were applied on both sides of each leave sample to keep it moist (Guo et al., 2015). ChlF was measured each morning and night in 12-hour intervals five times. The illumination light intensity was set at 3,000 μmol photons m^−2^s^−1^.

Polynomial fitting and curvature analysis were executed in Matlab (Mathworks, Natick, MA, USA). Statistical analysis was also performed to detect statistical significances in SPSS (Armonk, NY, USA).

## 4. Results

### 4.1 Curve Fitting and Curvature Analysis

Examples of polynomial fitting to experimental data are shown in Fig. 2. Fig. 2a is for a tree (*Pittosporum tobira*) leaf and Fig. 2b is for a spinach leaf. For a total of 722 sets of OJIP data, the relative fitting error is less than 0.018%, which shows that polynomials can represent the OJIP patterns very well.

**Fig. 2.**
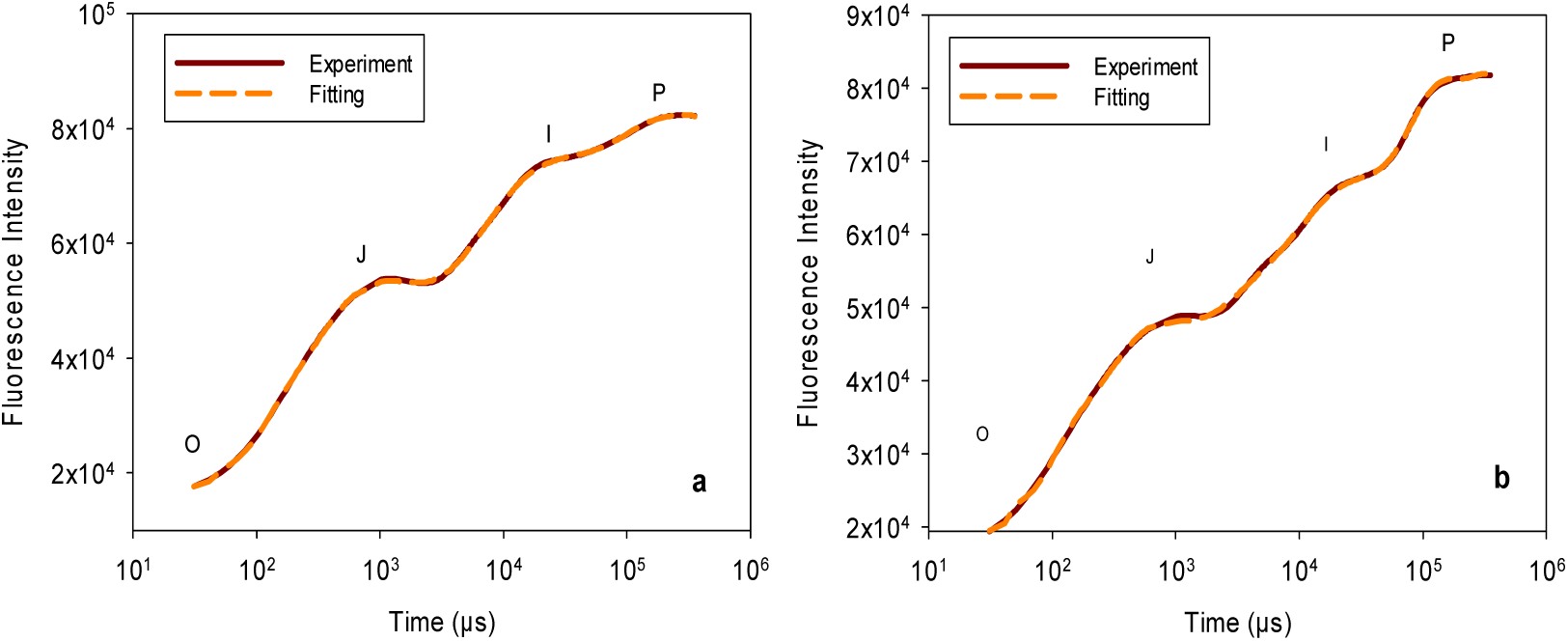
Comparison between experimental data and fitted polynomial. (a) A tree leaf (*Pittosporum tobira*), (b) A spinach leaf.

Figure 3 shows OJIP induction curves and the corresponding curvature values obtained with Eq. (4). Fig. 3a is the result for a tree (*Pittosporum tobira*) leaf and Fig. 3b for a spinach leaf. Obviously, there is a maximum value for each transition on the OJIP induction curve. These maximums provide quantitative information on OJIP characteristic transitions. The maxima around J and I appear in pairs: one corresponds to the upward transition and the other to the downward transition. Based on the descriptions of J and I in the literature, the maxima corresponding to the upward transitions are J or I. The peaks in the curvature are useful for quantitatively segmenting the OJIP curve into different phases.

**Fig. 3.**
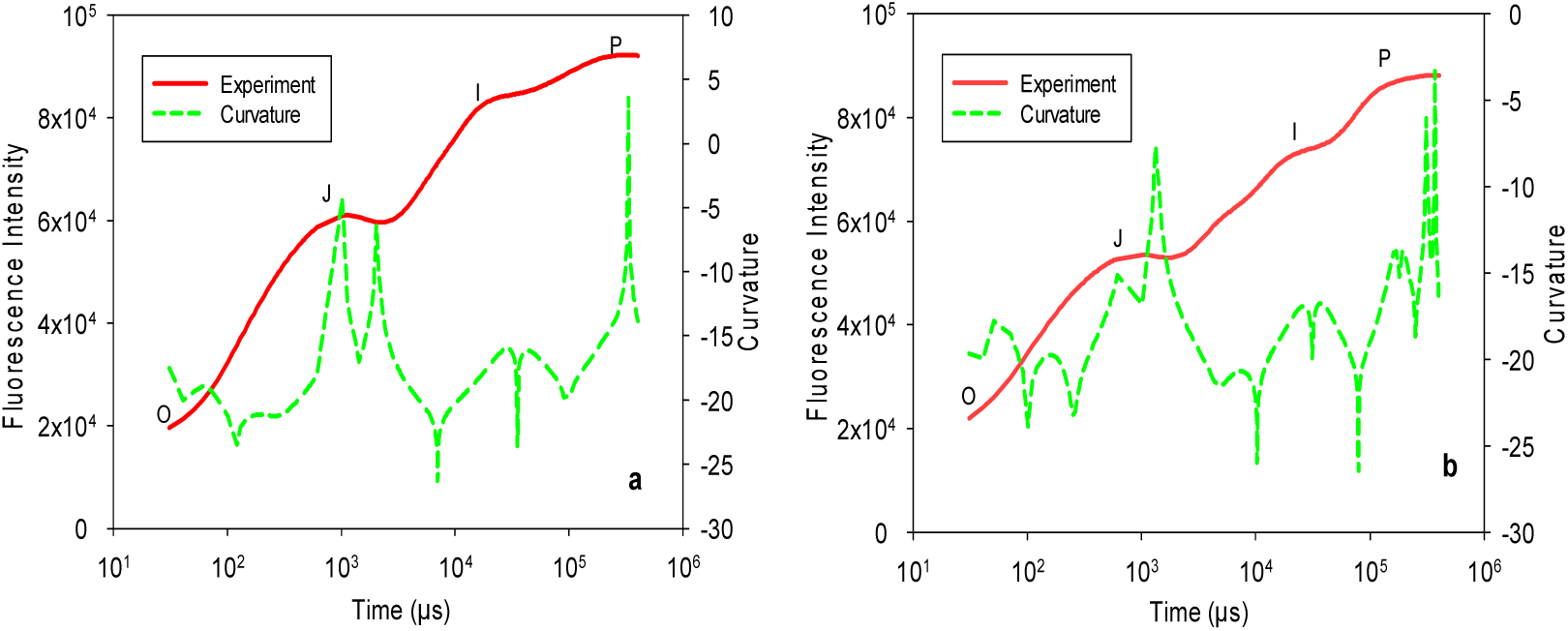
OJIP induction curve and corresponding curvature. (a) Result for a tree leaf (*Pittosporum tobira*); (b) Result for a spinach leaf.

As explained earlier, different samples may differ in physiological states, which may result in earlier or later times of occurrence for J and I. Even for the same leaf, if the environmental conditions such as temperature and illumination light intensity are changed, the J and I transitions may appear earlier or later. For a given leaf, it is difficult to determine what the optimal illumination light is before experiments. During experiments, light illumination changes the dark-adaptation status and thus it is difficult to try different experimental conditions. This causes errors in determining J and I transitions if a fixed time is used in ChlF meter protocols. Figures 4a and 4b show comparisons of the J and I transitions determined at the default times or by the curvature analysis. Figure 4a is for the leaf of *Cercis chinensis* at various temperatures and Fig. 4b is the result for *Cercis chinensis* at various light intensities. It is clear from Figures 4a and 4b that the curvature analysis method could track the transitions adaptively to always locate J and I at the inflection points as desired when the illumination light intensity and environmental temperatures change, whereas the traditional method of a fixed time misses the inflection points most times and by variable amounts. For all the measured *Cercis chinensis* samples, the time of occurrence for J (*t*_*j*_) varied from 1,293.41 μs to 2,127.90 μs and that for I (*t*_*i*_) varied from 30,734.79 μs to 47,941.69 μs. The default values in the instrumental protocol for J and I, however, are fixed at 2,012 μs and 30,321 μs, respectively. Distributions of the *t*_*j*_ and *t*_*i*_ determined by the developed method are shown in Figures 4c and 4d, respectively, which show considerable ranges of variation and thus inadequacy of using fixed times.

**Fig. 4.**
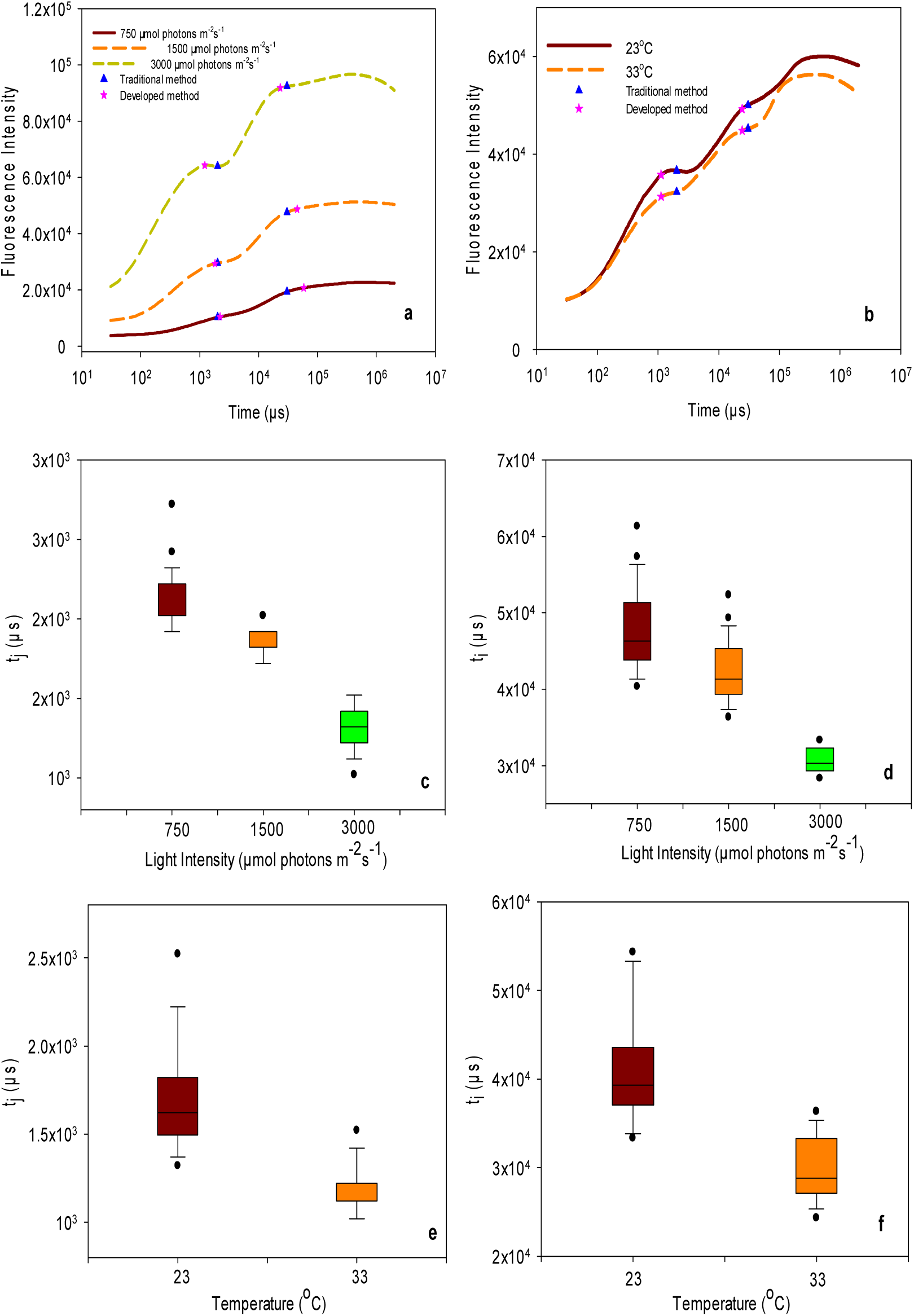
Comparisons of the fixed J and I transition points in instrumental protocol and those determined by the curvature analysis method. (a) Results for different light intensities (750, 1,500, and 3,000 μmol photons m^−1^ S^−1^), (b) Results for different temperatures, (c) Distribution of *t*_*j*_ for different light intensities, (d) Distribution of *t*_*i*_ for different light intensities, (e) Distribution of *t*_*j*_ for different temperatures, (f) Distribution of *ti* for different temperatures.

### 4.2 Differences in the times of occurrence and F_j_ and F_j_ values from different plants obtained by the developed method or the default ChlF meter protocol

The p-values (Appendix A) were examined for comparing the J and I times of occurrence and ChlF intensities obtained through the traditional method (*F*_*j*_, *T*_*j*_, *F*_*i*_, *T*_*i*_) and those by the developed method (*f*_*j*_, *t*_*j*_, *f*_*i*_, *t*_*i*_). The results reveal that the *F*_*j*_ and *f*_*j*_, *T*_*j*_ and *t*_*j*_ are statistically different for all the five types of plant leaves and that *F*_*i*_ and *f*_*i*_ and *t*_*j*_ are statistically different for some of the five types of plant leaves.

To show the effects of plant type, the ChlF variables obtained by the traditional method and those by the developed method were subject to t-tests (Appendix B). It should be mentioned that *T*_*j*_ and *T*_*i*_ were fixed and thus were not subject to the analyses. The results shown in Appendix B indicate that the measurements by the two methods differed significantly for the plant species in most cases. It is important to note that *t*_*j*_ is significantly different for all the five species and yet it is taken as a single fixed value in the traditional method.

From this set of experiments, it is clear that the proposed approach gave more consistent differences among different plant types. *t*_*j*_, in particular, consistently differed among all five plant types, whereas the traditional method uses a fixed value.

### 4.3 Differences in the times of occurrence and F_j_ and F_i_ values under varied light intensities obtained by the developed method or by the default ChlF meter protocol

Appendix C shows the means of ChlF variables determined by the default ChlF meter protocol or the developed method for the four types of leaves under different light intensities. Appendix D shows the *p* values of statistical comparison between the two methods under different light intensities, which reveal that most of the ChlF variables are statistically different. With increasing light intensity, the J and I points apparently moved to the left in general as revealed in Figure 4a and the *t*_*j*_ values shown in Appendix C. The times of occurrence for J and I varied significantly among different plant types and light intensities. However, in traditional instrumental protocol, *t*_*j*_ and *t*_*i*_ usually were set at fixed values. This unavoidably leads to discrepancies.

### 4.4 Differences in the times of occurrence and F_j_ and F_i_ values in various temperatures obtained by the developed method or by the default ChlF meter protocol

Appendix E shows the mean values of ChlF variables determined by the developed method or by the traditional method in room temperature (23°C) or an elevated temperature (33°C) and Appendix F shows the statistical comparison. Clearly, temperature has a significant influence on the occurring times of J and I transitions as revealed in Appendix D and Figure 4b. The developed method could automatically track the J and I transitions according to the curvature of the OJIP induction. Appendix E also reveals that lower temperature may result in higher values for *t*_*i*_ and *t*_*j*_. In this set of experiments, the J and I points apparently moved to the left and the obtained *t*_*j*_ and *t*_*i*_ values increased with the increments of temperature.

### 4.5 Differences in the times of occurrence and F_j_ and F_i_ values under different detachment and chilling stress obtained by the developed method or by the default ChlF meter protocol

To compare the differentiation abilities between the developed method and the default ChlF meter protocol, measurements from spinach leaves in the fourth set of experiments were used for statistical analysis and the results are shown in Appendix G. The leaves measured were subjected to five levels of detachment duration and chilling stress. In this group of experiments, the J and I points apparently moved to the right with the increase of duration of detachment and chilling stress. It can be easily found that all the statistically significant differences detected by the traditional method were detected by the corresponding developed method. However, the developed method especially *t*_*j*_ and *t*_*i*_, differentiated more group differences than the traditional method. *t*_*j*_ and *t*_*i*_ are unjustifiably set as constants in the traditional method.

## 5. Discussion

Differences in plant physiological status and experimental conditions such as illumination intensity and temperature will lead to significant changes in the shape of the OJIP induction curve and shift the J and I points either left or right. This was well demonstrated by the experiments in this work. A standard instrumental protocol uses fixed occurring times for the J and I transitions, which may catch the right J and I transition times for a specific plant type and under standardized conditions but may miss the inflection points by large margins under other circumstances as shown in Figure 4. This will lead to inconsistencies in measurement and introduce additional variations, which could reduce the usefulness of ChlF measurements.

The developed method based on curvature analysis can automatically locate J and I when differences in plant physiological status and environmental conditions shift the inflection points. ChlF measurements by the proposed method show better consistency and differentiation capabilities than those by the default instrument protocol. For example, *f*_*j*_ obtained by developed method showed consistent differences between the 2^nd^ morning and the 3^rd^ night for chilled spinach while the traditional method could not detect the difference.

Statistical comparison between the two methods under different light intensities, different temperatures and different plants show that there are statistically significant differences in most of the ChlF variables. The experimental results by the new method are consistent with what is known. For example, photosynthesis will speed up when the light intensity increases and faster reaction rates will lead to earlier J and I transitions, as shown by the new method.

The experimental results also show that *t*_*j*_ and *t*_*i*_ are very useful ChlF variables. *t*_*j*_ can clearly differentiate five types of leaves but in the standard method it is set as a fixed value. *t*_*i*_ and *t*_*j*_ can also be used to distinguish plants in different light intensities and different temperatures. While one may argue that the curvature method changes what have been conventionally understood as the J or I points, these new characterizations could be labeled differently and used because their differentiation capabilities as demonstrated in this work. Furthermore, this work indicates that other mathematically-definable features of the ChlF signal could prove useful as well.

## 6. Conclusion

PSII chlorophyll fluorescence characteristics have been widely used to sense plant physiological status and environmental changes. Differences in plant and experimental conditions will unavoidably result in different transition times. The method developed in this work can automatically detect and track the transitions. The measured characteristic fluorescence levels showed better differentiation abilities than those determined by the standard ChlF meter protocol method. The transition times determined show strong differentiation abilities for different plant species under different experimental conditions.

## Acknowledgements

This project is partially supported by National Natural Science Foundation of China (No: 31771680), the Fundamental Research Funds for the Central Universities of China (No: JUSRP51730A), the Modern Agriculture Funds of Jiangsu Province (No: BE2015310; No.SXGC[2017]210), the New Agricultural Engineering of Jiangsu Province (No.SXGC[2016]106), the 111 Project (B1208) and the Research Funds for New Faculty in Jiangnan University.

## References

Bradbury, M., & Baker, N. R. (1984). A quantitative determination of photochemical and non-photochemical quenching during the slow phase of the chlorophyll fluorescence induction curve of bean leaves. Biochimica et Biophysica Acta (BBA)-Bioenergetics, 765(3), 275–281.

Butler, W. L. (1978). Energy distribution in the photochemical apparatus of photosynthesis. Annual Review of Plant Physiology, 29(1), 345–378.

Coombs, J., Hall, D. O., & Long, S. P. (Eds.). (2014). Techniques in bioproductivity and photosynthesis: pergamon international library of science, technology, engineering and social studies. Elsevier.

Maxwell, K., & Johnson, G. N. (2000). Chlorophyll fluorescence—a practical guide. Journal of experimental botany, 51(345), 659–668.

D’Ambrosio, N., Arena, C., & De Santo, A. V. (2006). Temperature response of photosynthesis, excitation energy dissipation and alternative electron sinks to carbon assimilation in Beta vulgaris L. Environmental and experimental botany, 55(3), 248–257.

Gamon, J. A., Field, C. B., Bilger, W., Björkman, O., Fredeen, A. L., & Peñuelas, J. (1990). Remote sensing of the xanthophyll cycle and chlorophyll fluorescence in sunflower leaves and canopies. Oecologia, 85(1), 1–7.

Goltsev, V., Zaharieva, I., Lambrev, P., Yordanov, I., Strasser, R., 2003. Simultaneous analysis of prompt and delayed chlorophyll a fluorescence in leaves during the induction period of dark to light adaptation. J. Theor. Biol. 225, 171–183.

Govindjee, Sixty-three years since Kautsky: chlorophyll a fluorescence, Aust. J. Plant Physiol. 22 (1995) 131–160.

Guo, Y., Zhou, Y., & Tan, J. (2015). Wavelet analysis of pulse-amplitude-modulated chlorophyll fluorescence for differentiation of plant samples. Journal of theoretical biology, 370, 116–120.

Guo, Y., & Tan, J. (2015). Recent advances in the application of chlorophyll a fluorescence from photosystem II. Photochemistry and photobiology, 91(1), 1–14.

Kooten, O., & Snel, J. F. (1990). The use of chlorophyll fluorescence nomenclature in plant stress physiology. Photosynthesis research, 25(3), 147–150.

Papageorgiou, G. C., Tsimilli-Michael, M., & Stamatakis, K. (2007). The fast and slow kinetics of chlorophyll a fluorescence induction in plants, algae and cyanobacteria: a viewpoint. Photosynthesis research, 94(2-3), 275–290.

Krause, G. H., & Weis, E. (1991). Chlorophyll fluorescence and photosynthesis: the basics. Annual review of plant biology, 42(1), 313–349.

Koyro, H. W. (2006). Effect of salinity on growth, photosynthesis, water relations and solute composition of the potential cash crop halophyte Plantago coronopus (L.). Environmental and Experimental Botany, 56(2), 136–146.

Lavergne, J., & Trissl, H. W. (1995). Theory of fluorescence induction in photosystem II: derivation of analytical expressions in a model including exciton-radical-pair equilibrium and restricted energy transfer between photosynthetic units. Biophysical Journal, 68(6), 2474–2492.

Lubitz, W., Reijerse, E. J., & Messinger, J. (2008). Solar water-splitting into H 2 and O 2: design principles of photosystem II and hydrogenases. Energy & Environmental Science, 1(1), 15–31.

Mathur, S., Jajoo, A., Mehta, P., & Bharti, S. (2011). Analysis of elevated temperature induced inhibition of photosystem II using chlorophyll a fluorescence induction kinetics in wheat leaves (Triticum aestivum). Plant Biology, 13(1), 1–6.

Mohammed, G. H., Binder, W. D., & Gillies, S. L. (1995). Chlorophyll fluorescence: a review of its practical forestry applications and instrumentation. Scandinavian Journal of Forest Research, 10(1-4), 383–410.

Ogaya, R., Penuelas, J., Asensio, D., & Llusià, J. (2011). Chlorophyll fluorescence responses to temperature and water availability in two co-dominant Mediterranean shrub and tree species in a long-term field experiment simulating climate change. Environmental and Experimental Botany, 73, 89–93.

Rodriguez, M., Greenbaum, E., 2009. Detection limits for real-time source water monitoring using indigenous freshwater microalgae. Water Environ. Res. 81 (11), 2363–2371.

Ruban, A. V., & Belgio, E. (2014). The relationship between maximum tolerated light intensity and photoprotective energy dissipation in the photosynthetic antenna: chloroplast gains and losses. Philosophical Transactions of the Royal Society of London B: Biological Sciences, 369(1640), 20130222.

Schansker, G., Tóth, S. Z., Holzwarth, A. R., & Garab, G. (2014). Chlorophyll a fluorescence: beyond the limits of the QA model. Photosynthesis research, 120(1-2), 43–58.

Stirbet, A. (2011). On the relation between the Kautsky effect (chlorophyll a fluorescence induction) and photosystem II: basics and applications of the OJIP fluorescence transient. Journal of Photochemistry and Photobiology B: Biology, 104(1-2), 236–257.

Strasser, R. J. (1992). The Fo and the OJIP fluorescence rise in higher plants and algae. In Regulation of Chloroplast Biogenesis (pp. 423–426). Springer, Boston, MA.

Strasserf, R. J., & Srivastava, A. (1995). Polyphasic chlorophyll a fluorescence transient in plants and cyanobacteria. Photochemistry and photobiology, 61(1), 32–42.

Stirbet, A., Strasser, B. J., & Strasser, R. J. (1998). ChlorophyllaFluorescence Induction in Higher Plants: Modelling and Numerical Simulation. Journal of theoretical biology, 193(1), 131–151.

Vredenberg, W.J., 2004. System analysis of photoelectrochemical control of chlorophyll fluorescence in terms of trapping models of Photosystem II: a challenging view. In: Papageorgiou, G.C., Govindjee (Eds.), Chlorophyll a Fluorescence: A Signature of Photosynthesis Advances in Photosynthesis and Respiration, 19.

Taiz, L., Zeiger, E., 2006. Plant Physiology. Sinauer Associates Inc., Sunderland, pp. 139–148.

Zhu, X. G., Baker, N. R., Ort, D. R., & Long, S. P. (2005). Chlorophyll a fluorescence induction kinetics in leaves predicted from a model describing each discrete step of excitation energy and electron transfer associated with photosystem II. Planta, 223(1), 114–133.

Zhu, S., Zhang, J., Tang, S., Qiao, C., Wang, L., Wang, H., … & Wang, X. (2012). Surface chemistry routes to modulate the photoluminescence of graphene quantum dots: From fluorescence mechanism to up-conversion bioimaging applications. Advanced Functional Materials, 22(22), 4732–4740.

Zivcak, M., Brestic, M., Olsovska, K., 2008. Application of photosynthetic parameters in the screening of wheat (Triticum aestivum L.) genotypes for improved drought and high temperature tolerance. In: Allen, J.F.,

